# Modeling habitat suitability of vulnerable Mollucan Babirusa *Babyrousa babyrussa* in small island of Buru, Indonesia

**DOI:** 10.1101/2021.07.28.454166

**Authors:** Andri Wibowo

## Abstract

Babirusa is a mammal belongs to Suidae family. This mammal belongs to *Babyrousa* genus is known endemic to Indonesia. Recently there are 3 species of *Babyrousa*, one species is *Babyrousa babyrussa*, listed as ‘Vulnerable’ on the IUCN Red List of Threatened Species. *Babyrousa* occurs in Indonesia on Sulawesi island, Togian, Sula islands, and Taliabu, Mangole and Buru islands in the Molucca regions. The Moluccan Babirusa is now restricted to upland forests and mountainous terrain. Then this study aims to assess the suitability of Buru island as habitat for Moluccan Babirusa. The suitability analysis was based on GIS analysis using 4 determinant environmental variables required by *B. babyrussa* species including NDVI, barren soil, elevation, and river network. The particular location was a Batabual landscape sizing 292.60 km^2^ located in the east parts of Buru island. Based on NDVI, less vegetation covers were observed in north and east parts of Batabual. In contrast, NDVI values were higher in the central, west, and south indicating that half of the landscape was covered by intact primary forest. Batabual has a hilly landscape surrounded by coastal areas with hilly areas were observed in the south and west. There were 2 river streams in the east parts. The analysis has divided the landscape into several parts based on the habitat suitability levels. About one third of the landscape located in north was considered less and not suitable for Babirusa. Some areas in west were also considered moderate to be inhabited by Babirusa considering a presence of hilly landscape that may limit the vertical distribution of Babirusa. Most suitable habitats were estimated in central parts of the landscape spanning to the south. South parts of the landscape were characterized by high NDVI values and forest covers then these areas were considered as suitable habitats for Babirusa. The size of this suitable habitat was estimated around 188.62 km^2^ or more than half of the size (64.46%) of Batabual landscape.

## Introduction

In Indonesia, *Babyrousa* genus and its common name is Babirusa means “pig deer”; recognized from its featured four curled tusks, present only in the male. There are 3 species of *Babyrousa*, one species is *Babyrousa babyrussa*, listed as ‘Vulnerable’ on the IUCN Red List of Threatened Species. It occurs in Indonesia on Sulawesi island, Togian, Sula islands, and Taliabu, Mangole and Buru islands in the Molucca regions, and is confined to primary rainforest. Those islands have different species of Babirusa with *B. celebensis* for Sulawesi, *B. togeanensis* for Togian island, and *B. babyrussa* for Taliabu, Mangole and Buru islands. The Moluccan Babirusa is now restricted to upland forests and mountainous terrain, away from areas inhabited by humans.

Babirusa is gravely threatened by destruction of its rainforest habitat due to largescale commercial logging and agricultural conversion. There is also some illegal poaching of this species for its meat by local village communities. Babirusa has a low reproductive rate, and has only 1 to 2 piglets per year, which coupled with its highly restricted global range makes it vulnerable to rapid decline from any losses in its numbers.

Research on Babirusa especially the ecology and habitat of *Babyrousa babyrussa* is still limited currently. Data of Babirusa in particular islands are still lacking while the populations of this mammal species is declining. Then this paper aims to model the habitat suitability of vulnerable *Babyrousa babyrussa* in small island of Buru, Indonesia.

## Methodology

### Study area

The location was a Buru island part of Molluca ecoregion. This island has a size of 8,473 km^2^ with shore line length of 43.7 km. The highest point was 2,736 m located in the west part of the island and observed in mount Kalatmada. The landscape of Buru island was a combination of low land and mountainous region with hilly landscape (Cipta Karya 2011). Buru island still has a significant primary rainforest covers up to 59.98%. The rest land covers consisting of secondary forest (0.51%), mangrove (0.9%), and peat (0.06%). The intact land covers of Buru island have been converted into plantation (1.66%), swidden agriculture, (1.41%), and paddy field (1.82%). Settlements were mostly located in the coasts (0.41%) and near the roads. Particular study area was a Batabual (Figure 1) landscape sizing 292.60 km^2^ in the east parts of the island. This landscape has a hilly and forested landscape and intersected with several rivers in the west coast.

**Figure 1.**
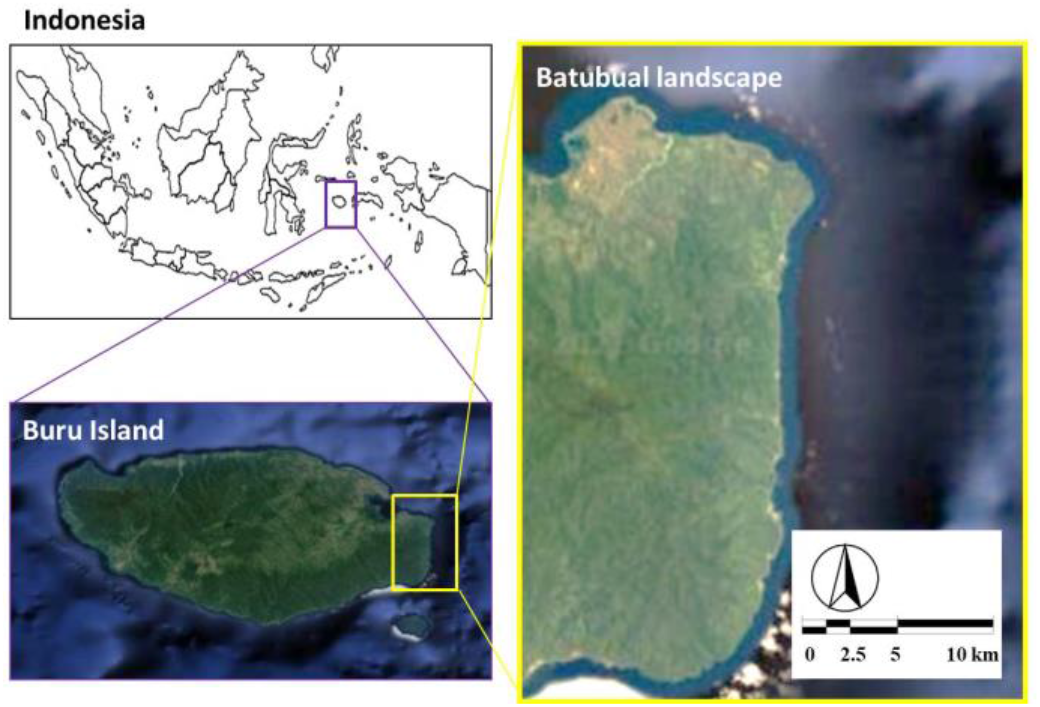
Study area in Batabual landscape, Buru island, Indonesia

### Habitat suitability analyses

The Babirusa habitat analysis was based on Geographical Information System (GIS) analysis. Within this GIS, the evaluated variables were related to the environmental variables that affect the presences of Babirusa. Those variables were including NDVI, barren soil, elevation, and rivers. NDVI represented the quality of vegetation condition that is related to the habitat and diet of Babirusa. Barren soil represented the magnitude of anthropogenic effects in the form of deforestation in reducing Babirusa habitats. Babirusa has specific elevation then landscapes with elevation of 0-1500 were selected. Babirusa is mammalian species that requires water sources then landscapes near waterways and rivers were prioritized.

### NDVI

NDVI is Normalized Difference Vegetation Index and NDVI of Batabual landscape was measured using method following Philiani et al. (2016), Kawamuna et al. (2017), and Sukojo and Arindi (2019). The NDVI is a simple graphical indicator used to analyze remote sensing measurements, often from a space satellite platform, assessing whether or not the target being observed contains live green vegetation. NDVI analyses wave length of satellite image retrieved from Landsat 8 containing vegetation image and in this study was forest covers. This measurement is possible since cell structure of the vegetation leaves strongly reflects near-infrared light wave length ranges from 0.7 to 1.1 µm. The calculation of NDVI for each pixel of vegetation pixel was as follows:

NDVI = near invisible red wave length – red wave length / near invisible red wave length + red wave length

The NDVI was denoted as a range from 0 (no vegetation) to 1(high vegetation density). The NDVI values then overlayed and mapped into Batabual landscape using GIS. The forest covers then categorized and classified by using NDVI as follows:

‐ if 0 < NDVI < 0.3 then forest covers < 50% (brown color)
‐ if 0.31 < NDVI < 0.4 then forest covers are 50 – 69% (pale brown color)
‐ if 0.41 < NDVI < 1.0 then forest covers are 70 – 100% (green color)

### Barren land

The habitat suitability analysis was measured using habitat characteristic that either favors or hinders the presence of the species. Presence of barren land representing deforestation will be avoided by the Babirusa. Barren land in this study was using method by Diek et al. (2017) and Zhao and Chen (2005) using satellite imagery and remote sensing. The proposed barren land (BI) equation was:

BI = (short wave infra red + red) – (near invisible red + blue) / (short wave infra red + red) + (near invisible red + blue).

The BI was denoted as a range from 0 (no barren land) to 1(high barren land). The BI values then overlayed and mapped into Batabual landscape using GIS. The barren lands then categorized and classified by using BI as follows:

‐ if 0 < BI < 0.3 then barren land covers < 50% (green color)
‐ if 0.31 < BI < 0.4 then barren land covers are 50 – 69% (green to red color)
‐ if 0.41 < BI < 1.0 then barren land covers are 70 – 100% (red color)

### Elevation

Elevation analysis was using the Digital Elevation Model (DEM) (Cheyne et al., 2016; Valerio et al. 2020). Elevation level of Batabual landscape was divided into several meter (m) classes above sea level. The DEM was represented as raster and denoted as m.

### River

Batabual landscape has several river ways. The rivers then were digitized and overlayed with the Batabual landscape. For Babirusa suitability analysis, the areas nearby rivers were having higher scores compared to the areas located far from the rivers.

## Results and Discussions

The measured variables that are related to the habitat suitability of Babirusa are available in Figure 2. Since it is a satellite imagery, then there were several cloud covers detected in the analysis. The vegetation covers as indicated by NDVI show that the vegetation covers across the landscape were varied. Low NDVIs were observed in the north and east of the landscape. Portion of landscape that has low NDVI in north was larger than in east. This indicates the less vegetation covers in north and east. In contrast, NDVI values were higher and reaching values close to maximum values were observed in the central, west, and south of the landscape. This indicates that half of the landscape was covered by intact primary forest.

**Figure 2.**
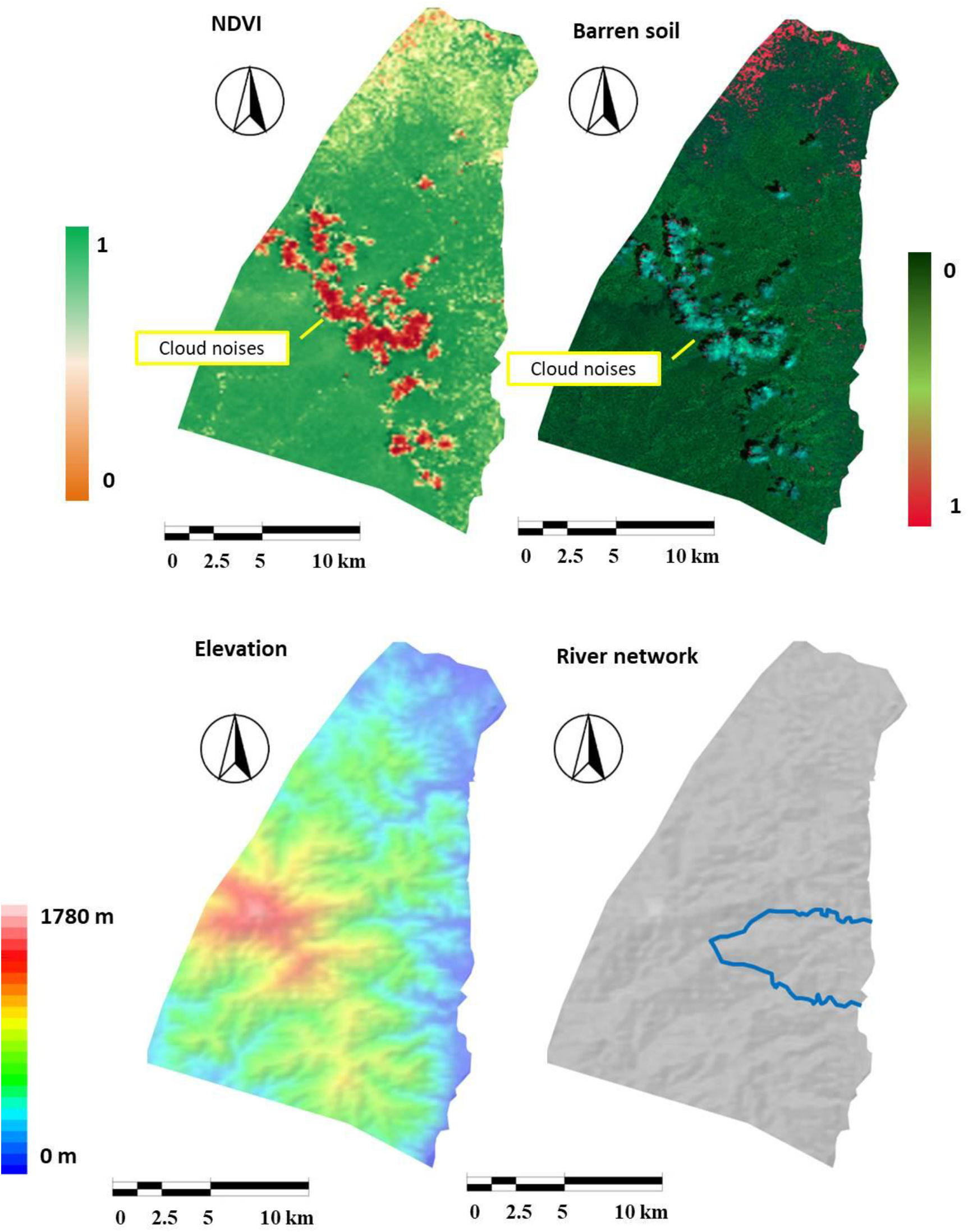
Environmental variables for Babirusa habitats in Batabual landscape, Buru island, Indonesia

High barren soil values were obviously detected from north west to north parts of the landscape. Some patches of landscape with high barren soil index were also detected in the east. This pattern of landscape fragment with high barren soil values was comparable to the fragment with low NDVI values. Batabual is a hilly landscape surrounded by coastal areas. The highest point was 1780 m above sea level. Hilly areas were observed in the south and west of Batabual. In contrast north and east parts of Batabual were dominated by low lands. A river network was observed in east part of the landscape. There were 2 river streams in the east parts. The upstream areas of the river were from the mountain. Then the east parts of the landscape were the river catchment areas of Batabual landscape.

The habitat suitability analysis was carried out by overlaying selected environmental variables. Those variables were having values that represent the suitability of those variables following Babirusa environmental requirements. The most suitable habitat was selected based on a portion of habitat that has the highest value. Figure 3 shows the estimated habitat for Babirusa in Batabual landscapes. The analysis has divided the landscape into several parts based on the suitability levels. About one third of the landscape located in north was considered less and not suitable for Babirusa. The reason for this considering that in this area, barren soils and low vegetation covers were detected in large portion. Some areas in west were also considered moderate to be inhabited by Babirusa considering a presence of hilly landscape that may limit the vertical distribution of Babirusa. Most suitable habitats were estimated in central parts of the landscape spanning to the south. Central parts were considered suitable since there were river networks considering Babirusa prefers wetlands. South parts of the landscape were characterized by high NDVI values and forest covers then these areas were considered as suitable habitats for Babirusa. The size of this suitable habitat was estimated around 188.62 km^2^or more than half of the size of Batabual landscape.

**Figure 3.**
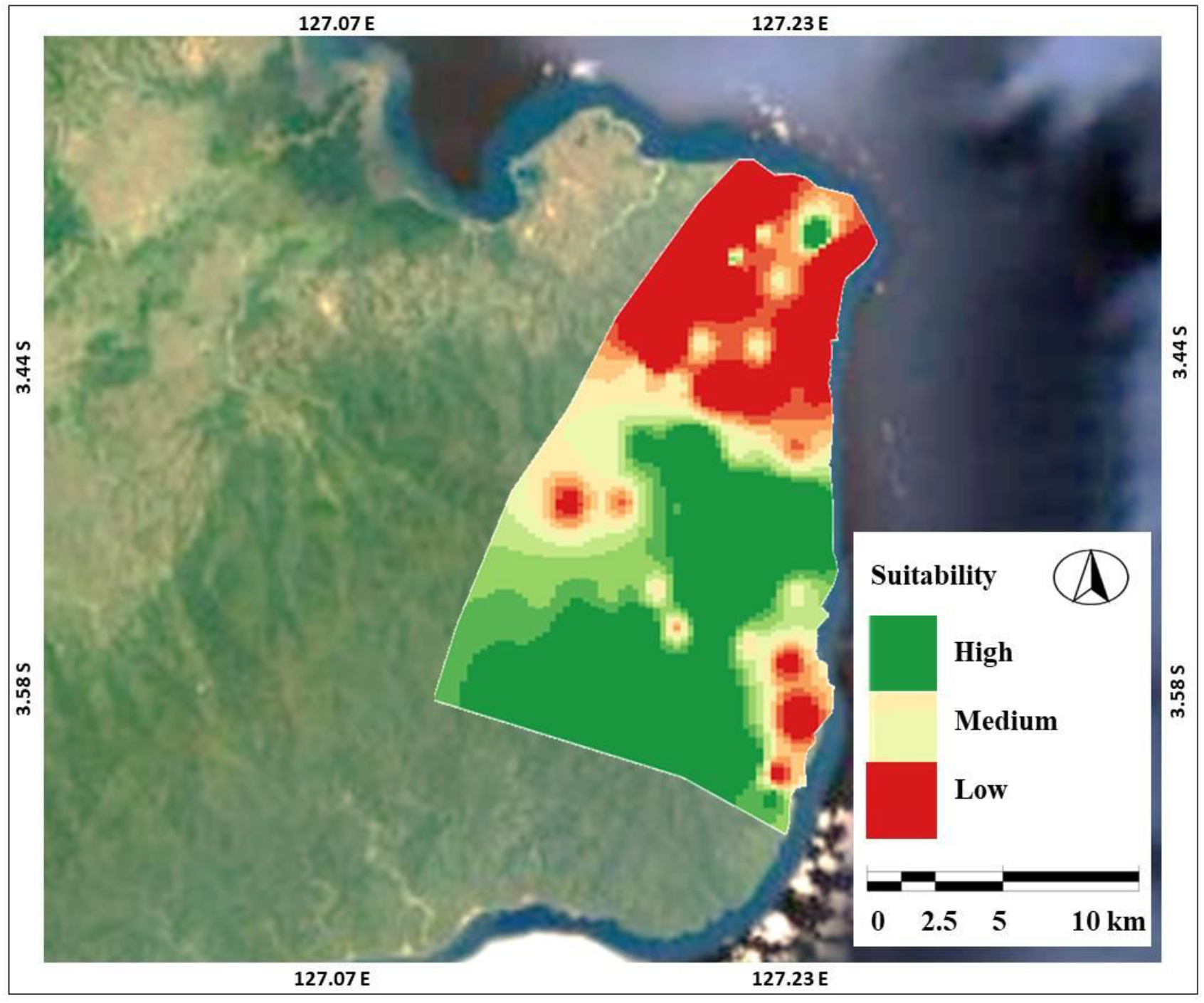
Estimated suitable habitats for Babirusa in Batabual landscape, Buru island, Indonesia

Selection of areas nearby river networks located in the east parts of Batabual considering the requirements of Babirusa. This Suid mammal like forest areas where there are rivers, swamps, and water inlets that allow it to get drinking water and wallow. The Babirusa also visits springs and salt-licks on a regular basis, in search of mineral salts that aid in digestion. Babirusa is often seen bathing (Anwarhadi et al. 2018) in puddles where the water is rather clean and not muddy. In summer it is often seen bathing in the river and this explain the selection of areas nearby river networks as the suitable habitats

This study is comparable to previous study by Rosyidy and Wibowo (2020) in estimating Babirusa’s suitable habitats in Gorontalo, parts of Sulawesi. The difference in this study was using river networks as determinant variables. Barren soil in here is a variable that can reduce suitability of measured variables. Barren soil is indicating the presence of anthropogenic activities. Some areas are considered not suitable for Babirusa habitat, that is areas and land use characterized by high intensity of human activities, including settlements, plantations, agriculture, rice fields, and fields. Whereas, a suitable area is forest and swamp, and occasionally hilly terrain far from human habitation. The Moluccan Babirusa is most often seen in the morning and late afternoon with an ability to run uphill quickly and an ability to swim well include often swimming in the sea to reach offshore islands. Then coastal areas should be considered and included as suitable habitats.

## Conclusion

GIS based suitable habitat assessment for medium and large mammals is important to support the species conservation. Previous work has been carried out for hippo species in Africa (Buruso, 2017) while similar researches are still lacking in Asia continent. South East Asia region mainly Indonesia archipelago is known with its high mammal diversity while the threats are also significant. Then this research has contributed significantly for mammals conservation mainly listed as vulnerable.

## References

Anwarhadi, N.Y., Labiro, E., Korja, I.N. 2018. Komposisi Vegetasi Habitat Babirusa (Babyrousa Babyrussa) Di Kawasan Hutan Pendidikan Universitas Tadulako Kecamatan Bolano Lambunu Kabupaten Parigi Moutong (in Bahasa). Jurnal Warta Rimba. 6(4).

Buruso, F. 2017. Habitat suitability analysis for hippopotamus (H. amphibious) using GIS and remote sensing in Lake Tana and its environs, Ethiopia. Environmental Systems Research. 6. 10.1186/s40068-017-0083-8.

Cheyne, S., Sastramidjaja, W., Muhalir Rayadin, Y., Macdonald, D. 2016. Mammalian communities as indicators of disturbance across Indonesian Borneo. Global Ecology and Conservation. 7. 157–173. 10.1016/j.gecco.2016.06.002.

Cipta Karya. 2011. Gambaran Umum Kondisi Kabupaten Pulau Buru (in Bahasa).

Diek, S., Fornallaz, F., Schaepman, M., Jong, R. 2017. Barest Pixel Composite for Agricultural Areas Using Landsat Time Series. Remote Sensing. 9. 1245. 10.3390/rs9121245.

Kawamuna, A.l., Suprayog, i A., Wijaya, A.P. 2017. Analisis Kesehatan Hutan Mangrove Berdasarkan Metode Klasifikasi NDVI Pada Citra Sentinel-2 (Studi Kasus : Teluk Pangpang Kabupaten Banyuwangi. Jurnal Geodesi Undip, 6(1): 277–284.

Philiani, I., Saputra, L., Harvianto, L., Muzaki, AA. 2016. Pemetaan Vegetasi Hutan Mangrove Menggunakan Metode Normalized Difference Vegetation Index (NDVI) Di Desa Arakan, Minahasa Selatan, Sulawesi Utara.

Rosyidy, M. & Wibowo, A. 2020. GIS-Based Spatial Model for Habitat Suitability of Babirusa (Babyrousa celebensis), in Gorontalo Province. 4. 10.7454/jglitrop.v4i1.77.

Sukojo, B.M., Arindi, Y.N. (2019). Analisa Perubahan Kerapatan Mangrove Berdasarkan Nilai Normalized Difference Vegetation Index Menggunakan Citra Landsat 8 (Studi Kasus: Pesisir Utara Surabaya). Geoid Journal of Geodesy and Geomatics 14(2).

Valerio, F., Ferreira, E., Godinho, S., Pita, R., Mira, A., Fernandes, N., Santos, S. 2020. Predicting Microhabitat Suitability for an Endangered Small Mammal Using Sentinel-2 Data. Remote Sensing. 12. 10.3390/rs12030562.

Zhao, H. & Chen, X.. 2005. Use of normalized difference bareness index in quickly mapping bare areas from TM/ETM+. International Geoscience and Remote Sensing Symposium (IGARSS). 3. 1666–1668.

